# Comparative Genomics of *Staphylococcus* Reveals Determinants of Speciation and Diversification of Antimicrobial Defense

**DOI:** 10.1101/277400

**Authors:** Rosanna Coates-Brown, Josephine Moran, Pisut Pongchaikul, Alistair Darby, Malcolm J. Horsburgh

## Abstract

The bacterial genus *Staphylococcus* comprises diverse species with most being described as colonizers of human and animal skin. A relational analysis of features that discriminate its species and contribute to niche adaptation and survival remains to be fully described. In this study, an interspecies, whole-genome comparative analysis of 21 *Staphylococcus* species was performed based on their orthologues. Three well-defined multi-species groups were identified: group A (including *aureus*/*epidermidis*); group B (including *saprophyticus*/*xylosus*) and group C (including *pseudintermedius*/*delphini*). The machine learning algorithm Random Forest was applied to identify variable orthologues that drive formation of the *Staphylococcus* species groups A-C. Orthologues driving staphylococcal infrageneric diversity comprised regulatory, metabolic and antimicrobial resistance proteins. Notably, the BraSR (NsaRS) two-component system (TCS) and its associated BraDE transporters that regulate antimicrobial resistance distinguish group A *Staphylococcus* species from others in the genus that lack the BraSR TCS. Divergence of BraSR and GraSR antimicrobial peptide survival TCS and their associated transporters was observed across the staphylococci, likely reflecting niche specific evolution of these TCS/transporters and their specificities for AMPs. Experimental evolution, with selection for resistance to the lantibiotic nisin, revealed multiple routes to resistance and differences in the selection outcomes of the BraSR- positive species *S. hominis* and *S. aureus*. Selection supported a role for GraSR in nisin survival responses of the BraSR-negative group B species *S. saprophyticus*. Our study reveals diversification of antimicrobial-sensing TCS across the staphylococci and hints at differential relationships between GraSR and BraSR in those species positive for both TCS.

**Importance:** The genus *Staphylococcus* includes species that are commensals and opportunist pathogens of humans and animals. Identifying the features that discriminate species of staphylococci is relevant to understanding niche selection and the structure of their microbiomes. Moreover, the determinants that structure the community are relevant for strategies to modify the frequency of individual species associated with dysbiosis and disease. In this study, we identify orthologous proteins that discriminate genomes of staphylococci. In particular, species restriction of a major antimicrobial survival system, BraSR (NsaRS), to a group of staphylococci dominated by those that can colonize human skin. The diversity of antimicrobial sensing loci was revealed by comparative analysis and experimental evolution with selection for nisin resistance identified the potential for variation in antimicrobial sensing in BraRS-encoding staphylococci. This study provides insights into staphylococcal species diversity.

## Introduction

### *Staphylococcus* species and genomics

The existence of taxonomically distinct species groups was first proposed for *Staphylococcus* based on differential DNA-DNA hybridization methods (1). These groups were supported by 16S rDNA sequence analysis of 38 taxa (2) and multilocus sequence data of around 60 species and subspecies (3).

A comparative analysis that utilized next generation genome sequencing data of staphylococci to probe phylogenetic relationships with 491 shared orthologues across 12 *Staphylococcus* species (4) proposed *S. pseudintermedius* then *S. carnosus* as basal lineages. Moreover, with ten species in their analysis being residents of human skin, the authors proposed that evolution selected for human adaptation after branching from *S. carnosus*. The relationships between the strains generated from shared orthologues were maintained using total gene content (4). However, in contrast to the conclusions of 16S rDNA and multilocus data (2,3) their analysis revealed discrete clustering of *Staphylococcus* species. No distinct clustering of *S. hominis* with *S. haemolyticus* was observed and *S. saprophyticus* was assigned to the *S. epidermidis* group of species (4). Currently, there is a knowledge gap in *Staphylococcus* species comparisons with a need to determine if this clustering of staphylococcal species is supported using whole genome data. Our findings here begin to close this gap.

## Two component systems

Prokaryotes are receptive to environmental stimuli through diverse sensory and transducing two component systems (TCS). These TCS archetypically comprise a sensor histidine kinase (HK) that spans the cell membrane to interact with the external environment. Stimulus perception causes conditional autophosphorylation that is relayed to an interacting response regulator (RR) to enable DNA-binding directed transcription modulation (5).

While TCS are widespread and diverse across prokaryotes, the intramembrane-sensing histidine kinases (IM-HK) are specific to the Firmicutes. This family of small HKs has a short, 25 amino acid linker region between each 400 amino acid transmembrane helix. *S. aureus* GraSR uses a IM-HK to regulate a global network responsible for resistance to antimicrobial peptides (AMPs). GraSR modulates the expression of DltABCD and MprF that in concert alter the *S. aureus* surface charge to evade electrostatic interaction-mediated targeting of cationic AMPs (6).

An orthologous TCS to GraSR described in *S. aureus* was concurrently designated BraSR and NsaSR by two different groups (7,8). Serial passage in sub-MIC concentrations of the lantibiotic nisin was shown to select increased nisin MIC due to a SNP in *nsaS* gene encoding sensor histidine kinase of NsaRS (nisin susceptibility-associated sensor regulator) (8). The TCS was separately designated BraSR (bacitracin resistance-associated sensor regulator) from the reduced MIC of bacitracin and nisin determined for the TCS gene mutant (7). BraR binding sites were revealed upstream of the ABC transporter genes *braDE* and *vraDE* that were not transcribed in the mutant but induced in the presence of bacitracin. The transporter BraDE contributes to the detection of nisin and bacitracin and subsequent signal transduction via BraSR, whereas VraDE is more directly involved in detoxification by efflux (7). Transcription of *braSR* is increased following exposure to multiple antibiotics, including ampicillin, phosphomycin and nisin. Inactivation of *braS* (*nsaS*) revealed differential transcription of 245 genes (9), revealing the TCS might report cell envelope stress to directly regulate biofilm formation, cellular transport and responses to anoxia.

In this study, a comparative genome analysis of 21 *Staphylococcus* species was performed based upon their orthologous gene content. Species groups were revealed and then interrogated using the Random Forest algorithm to identify group-contributing genes. The operon encoding the BraSR TCS was found to differentiate the *S. aureus/S. epidermidis* species group from other species groups. Experimental evolution of representative *braSR*-positive and -negative species with nisin selection identified differential selection of BraSR and GraSR to produce resistance to this AMP.

## Results and Discussion

### Analysis of orthologous gene content across the staphylococci

The orthologous gene content of 21 sequenced staphylococcal species’ genomes was determined using OrthoMCL to group orthologous genes (homologues separated by speciation) into clusters across the different species. The number of shared orthologous clusters between the different species’ genomes was then represented as a heatmap (Figure 1). The output from this analysis revealed the assembly of three major groups of species, each with high numbers of shared orthologous clusters. An associated cladogram supported three groups (groups A, B and C) when defined as containing three or more species (Figure 1). This supported previous reported groupings from 16S rDNA and multilocus analyses (2,3). Presented here, three species pairs showed a high degree of shared orthologous clusters of genes and branched together in the cladogram: *S. aureus*/*S. simiae*, *S. simulans*/*S. carnosus*, and *S. lentus/S. vitulinus*.

**Figure 1:**
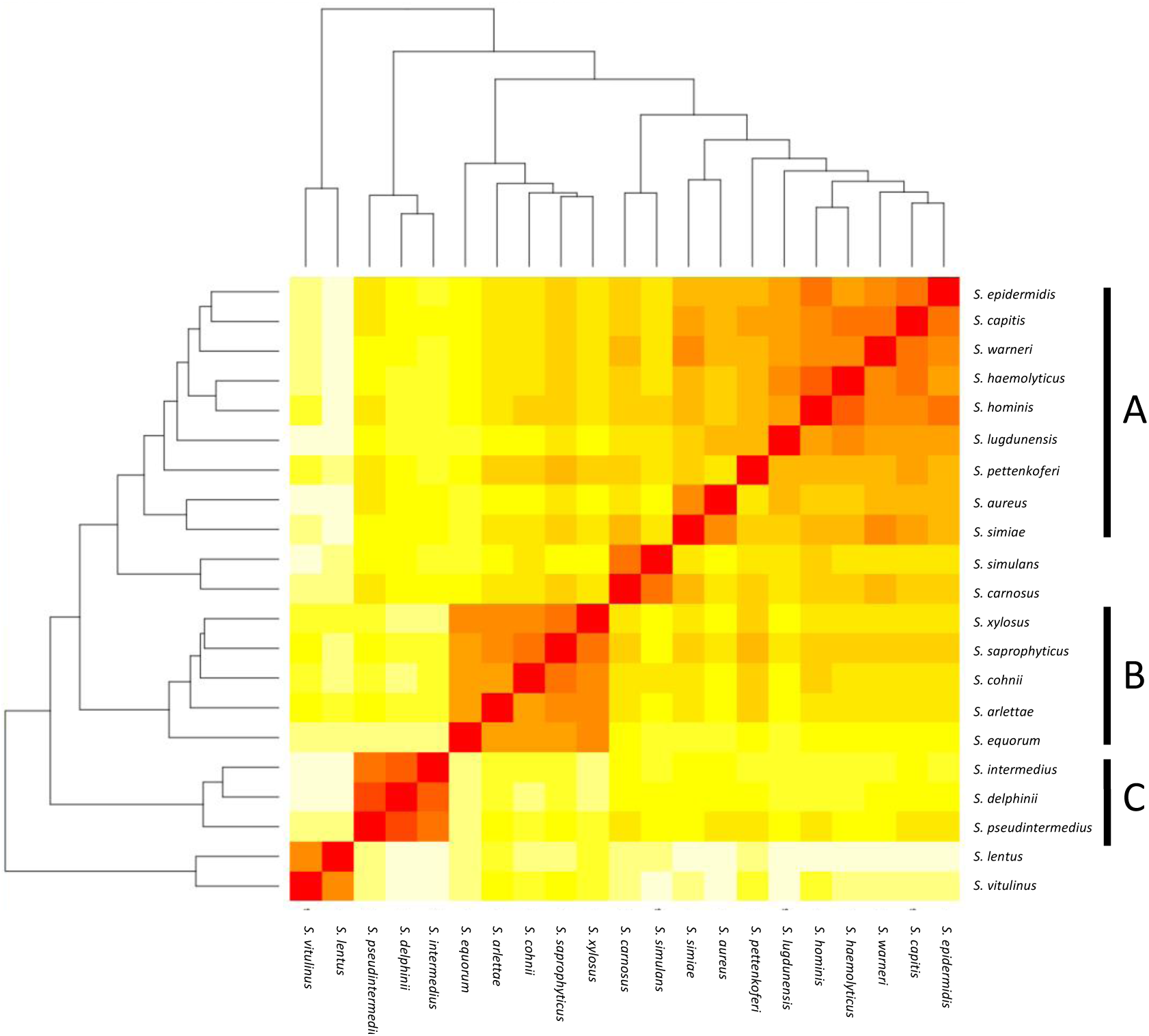
Heat map representation of shared orthologous proteins across *Staphylococcus* species. Presence is indicated using a color scale from red (highest number of shared clusters of orthologous proteins) to white (lowest number). Major groups of species observed in the analysis are highlighted as groups A-C

The largest and least well-defined, species group comprises *S. epidermidis*, *S. capitis*, *S. warneri*, *S. haemolyticus*, *S. hominis*, *S. lugdunensis*, *S. pettenkoferi*, *S. aureus* and *S. simiae* (Figure 1). Designated group A, it is dominated by species that colonize human skin (20,21). The likelihood of a strain-dependent effect structuring group A was investigated by substituting *S. epidermidis, S. hominis* and *S. aureus* strains (Supplementary File S1). Substituting these individual species with alternative strains and repeating the OrthoMCL analysis did not alter species groupings. Groups B and C were similarly unaffected by switching strains of *S. saprophyticus* and *S. pseudintermedius*, respectively. Within group A, *S. aureus* and *S. simiae* are most related in the cladogram; these species were proposed as members of the *S. aureus* group of staphylococci from gene content (2).

The smaller species group B comprises *S. equorum*, *S. arlettae*, *S. cohnii*, *S. saprophyticus* and *S. xylosus* (Figure 1). Though not universal, a frequent lifestyle identified in the group B species is human or animal host colonization; several species are associated with meat products and novobiocin resistance (22, 23) with commonalities in their cell wall composition (24).

Species group C comprises *S. pseudintermedius, S. delphini* and *S. intermedius* and this collective was previously designated the *S. intermedius* group (SIG); the species cause opportunistic infection of companion animals and equids (22). Emerging antibiotic resistance in the SIG species group is a clinical veterinary concern (25) and their routine speciation is complicated by their high degree of 16S rRNA locus sequence identity (26).

Genetic determinants directing the formation of species group A were tested in R using machine learning with the Random Forests algorithm for classification (27). This algorithm was used to identify variables, in this case OrthoMCL clusters, that contributed to formation of the groups, based on a forest of trees generated from these variables. OrthoMCL clusters representing each variable were determined and mapped back to respective genomes for each cluster and the PROKKA annotation of each protein coding sequence was verified using BLAST. Contributing variables were assigned for group A, based on the strain set described in Table 1, where permutations were used to verify the existence and reproducibility of species groups (Supplementary File S1).

**Table 1:** *Staphylococcus* species and strains included in OrthoMCL analysis. Genomes were sequenced for this study, as indicated, or retrieved from NCBI for analysis; genome integrity is indicated.

| **Species** | **Strain** | **Accession** | **Status (Reference)** |
| --- | --- | --- | --- |
| *S. arlettae* | CVD059 | ALWK01000000<br>(Uid175126) | Draft<br>(10) |
| *S. aureus* | Newman | AP009351<br>(Uid58839) | Complete<br>(11) |
| *S. capitis* | SK14 | ACFR01000000<br>(Uid55415) | Draft |
| *S. carnosus* | TM300 | NC012121<br>(Uid59401) | Complete<br>(12) |
| *S. cohnii* | ATCC29974 | LT963440 | Draft<br>(This study) |
| *S. delphini* | 8086 | CAIA00000000<br>(Uid199664) | Draft |
| *S. epidermidis* | ATCC_12228 | NC005008<br>(Uid57861) | Complete<br>(13) |
| *S. equorum* | Mu2 | CAJL01000000<br>(Uid169178) | Draft |
| *S. haemolyticus* | K8 | LT963441 | Draft<br>(This study) |
| *S. hominis* | J6 | LT963442 | Draft<br>(This study) |
| *S. intermedius* | NCTC_11048 | CAIB01000000<br>(Uid199665) | Draft |
| *S. lentus* | F1142 | AJXO01000000<br>(Uid200144) | Draft<br>(14) |
| *S. pettenkoferi* | VCU012 | AGUA00000000<br>(Uid180074) | Draft |
| *S. lugdunensis* | HKU09 | CP001837<br>(Uid46233) | Complete<br>(15) |
| *S. pseudintermedius* | HKU10 | Uid62125 | Complete<br>(16) |
| *S. saprophyticus* | ATCC_15305 | AP008934<br>(Uid58411) | Complete<br>(17) |
| *S. simiae* | CCM_7213 | AEUN00000000<br>(Uid77893) | Draft<br>(4) |
| *S. simulans* | ATCC 27848 | LT963435 | Draft<br>(This study) |
| *S. vitulinus* | F1028 | AJTR00000000<br>(Uid200114) | Draft<br>(18) |
| *S. warneri* | SG1 | CP003668<br>(Uid187059) | Complete<br>(19) |
| *S. xylosus* | ATCC29971 | LT963439 | Draft<br>(This study) |

### Clusters driving formation of group A species

The presence of 15 and absence of 9 OrthoMCL clusters collectively contribute to defining group A, with differing levels of support (Mean Decrease in Accuracy [MDA] values) (Table 2 & Supplementary File S2). Lower MDA values correspond with the incomplete presence of orthologous clusters within all species in a group. Several strongly supported orthologues contribute to group A (Table 2), notably the presence of four that are sequentially encoded in the genome as an operon (epi_02134 - epi_02137; MDA 3.2, 3.0, 2.6, 2.2, respectively). The latter cluster pair epi_02136/epi_02137 was annotated by PROKKA as a TCS sensor/regulator (Table 2 & Supplementary File S2) and shares ∼100% similarity with BraSR (SA2417/SA2418 of *S. aureus* N315), a TCS associated with resistance to AMPs nisin and bacitracin (7). The adjacent clusters encoded in the same operon (epi_02134, epi_02135) comprise the BraD/BraE ABC transporter subunits with 98% and 99% similarity with SA2415/ SA2416 of *S. aureus* N315, respectively (7). We demonstrate as a key finding of our analysis that the genomes of group A *Staphylococcus* species uniquely contain BraSR and BraDE.

**Table 2:** Proteins driving formation of species group. A. PROKKA annotation was found by mapping clusters from the variable importance analysis to the *S. epidermidis* genome in the case of ‘present’ clusters and the *S. saprophyticus* genome for ‘absent’ clusters. The PROKKA locus tag is indicated in brackets. BLAST homology was determined from searches against the NCBI BLAST database. %MDA is representative for the cluster across the randomForest analyses, where higher values indicate increased support.

| **Group A staphylococci** |  |  |
| --- | --- | --- |
| **PROKKA annotation** | **BLAST homology** | **MDA** |
| **Presence** |  |  |
| Putative cell wall associated hydrolase (epi_00542) | Hypothetical protein | 2.2 |
| Hypothetical protein (epi_02098) | Cell wall surface anchor protein | 2.7 |
| Hypothetical protein (epi_02108) | Hypothetical protein | 1.9 |
| FtsX like permease family protein (epi_02134) | ABC transporter permease | 3.2 |
| Macrolide export ATP binding/ permease protein MacB (epi_02135) | Bacteriocin ABC transporter ATP-binding protein | 3.0 |
| Sensor histidine kinase GraS (epi_02136) | TCS histidine kinase | 2.6 |
| Glycopeptide resistance associated protein R (epi_02137) | TCS transcriptional regulator | 2.2 |
| **Absence** |  |  |
| Succinate semialdehyde dehydrogenase NADP<sup>+</sup> (sap_00201) | Succinate-semialdehyde dehydrogenase | 2.9 |
| Putative membrane protein putative toxin regulator (sap_00203) | PTS sugar transporter subunit IIC | 2.7 |
| Putative multidrug resistance ABC transporter ATP binding/permease protein YheI (sap_00398) | Multidrug ABC transporter ATP-binding protein | 3.3 |
| Putative multidrug resistance ABC transporter ATP binding/permease protein YheH (sap_00399) | Multidrug ABC transporter ATP-binding protein | 1.4 |
| L-lactate utilization operon repressor (sap_00760) | Transcriptional regulator | 2.7 |
| Glutamate aspartate carrier protein (sap_01003) | Sodium:dicarboxylate symporter | 2.7 |

The presence of orthologue epi_00542 (MDA 2.2; Table 2 & Supplementary file S2) contributes to species group A, with support that the protein functions as a putative cell wall hydrolase from the Nlp-P60 family hydrolase domain that is associated with hydrolysis of peptidoglycan. Also, contributing to defining group A are the absences of two orthologue clusters (sap_00398; MDA 3.3 and sap_00399; MDA 1.4; Table 2 & S2) that are annotated as multidrug ABC transporters. A range of cytotoxic molecules are mobilized across the cell membrane by multidrug ABC transporters where certain families of these can also act as sensors (5, 28). Across staphylococcal groups, differential repertoires of ABC transporters associated with antimicrobial survival are consistent with the importance of community competition in species evolution.

Sequence variation of the NADP-dependent succinate semialdehyde dehydrogenase (SSADH) between group A staphylococci versus groups B and C was identified by the association of cluster sap_00201 (MDA 2.9, Table 2 & Supplementary File S2) with group A species; this variation might be allied to differences in glutamate metabolism across the genus. Glutamate is involved in multiple metabolic processes and bacterial glutamate dehydrogenase catabolizes glutamate, which contributes to acid tolerance.NADP-SSADH catalyzes catabolism of γ-aminobutyrate, a product of glutamate dehydrogenase activity (29); this pathway is oxidative stress sensitive owing to the catalytic cysteine residue of SSADH.

### Clusters driving formation of group B and C species

The size of species input groups B and C (Figure 1) used here limits use of the random forest algorithm and a broader species comparison of staphylococci could be considered in future. Consequently, a similar species-defined analysis of groups B and C was not pursued beyond a cursory view. Of note, however, a unique discriminating orthologue of group B was identified that supports broader species analyses. This orthologue is annotated as squalene synthase (SQS), a farnesyl diphosphate:farnesyl transferase (xylosus_00489; Supplementary File S3). SQS catalyzes an alternative pathway for the biosynthesis of carotenoid pigments (30). In *S. aureus*, biosynthesis of the carotenoid staphyloxanthin is catalyzed by the dehydrosqualene synthase CrtM, which converts farnesyl diphosphate (FPP) to dehydrosqualene (31). A recent study has determined that combining the activities of SQS and dehydrosqualene desaturase (CrtN) was sufficient to synthesize the carotenoid pigment staphyloxanthin by combining *E. coli* and *S. aureus* enzymes, respectively (31). Experimentation is required to determine catalytic specificity of the putative *Staphylococcus* group B-associated SQS. The *S. saprophyticus* protein has 13.24% similarity with *S. aureus* CrtM and 28% similarity with *Methylomonas* SQS. In support of differential substrate inputs, group B staphylococci such as *S. xylosus* and *S. arlettae* also encode CrtM in addition to CrtPQN necessary for staphyloxanthin biosynthesis.

### Diversity of cationic AMP survival loci across the staphylococci

The described comparative genomic analysis revealed that while BraSR TCS is restricted to group A species of staphylococci, the GraSR TCS is distributed across all species groups. Supporting predictions from the Random Forest analysis, low sequence identity of BraR/BraS with GraR/GraS was confirmed. BraR mean sequence identity with GraR of group A (44%) and group B/C species (40%) was greater than that of BraS compared with GraS of group A and groups B/C (mean ∼30% and ∼26%, respectively) (Table 3).

**Table 3:**
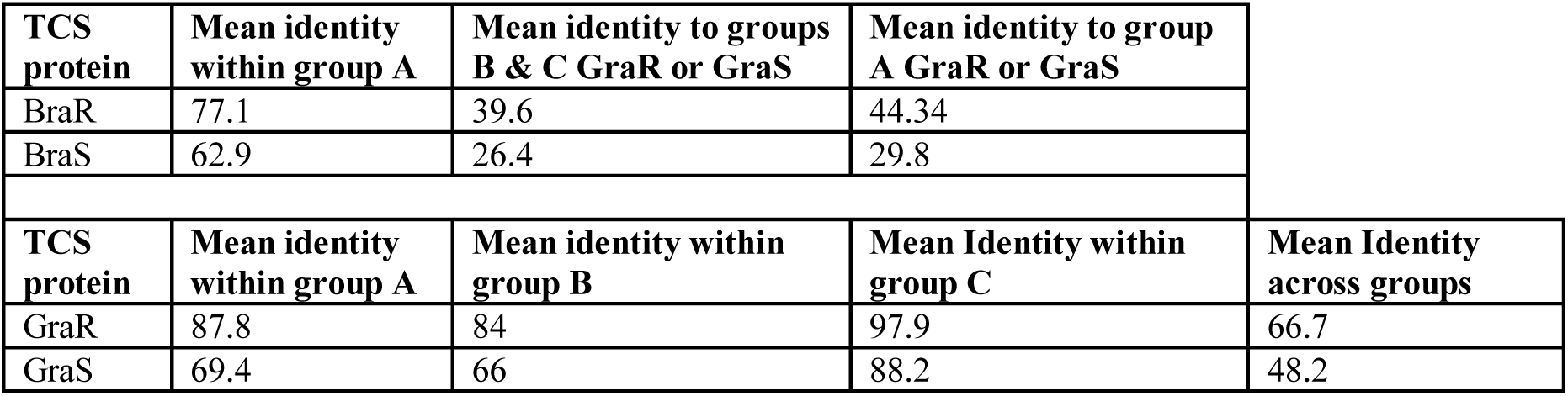
Comparative sequence identity of the BraRS and GraRS TCS across species groups A-C. Mean identity values of BraR and BraS within species group A and values for BraR and BraS with GraR and GraS of species group A or B/C. Mean identity values of GraR and GraS within and between species groups A, B and C. Sequence identity was calculated from multiple sequence alignments of all protein sequences of species indicated in Table 1.

High mean sequence identity (84-98%) of GraR regulator protein occurs within each of the three species groups (Table 3) with divergence of GraR between species groups identified by lower mean sequence identity (67%). GraS sensor histidine kinase was less conserved within species groups A (mean 69%) and B (mean 66%), compared with GraS of species group C that shared greatest mean sequence identity (88%), albeit that group C defined here is a small, related species set. Both BraS and GraS sensor proteins have lower sequence conservation across staphylococci than BraR and GraR (Table 3). The reduced divergence of these response regulators might reflect their relative isolation from selection by the external environment and differential stimuli.

Responses to cationic AMPs in the staphylococci are complex (32) and ligand specificity could account for species divergence of GraSR and BraSR TCS. This evolutionary outcome could be explained with strong selection pressure driven by ubiquity and diversity of cAMPs in staphylococcal niches. One intrigue in our analysis is the absence of GraSR and presence of only BraSR TCS in the group A species, *S. pettenkoferi*, with the sole related sensor protein having a mean sequence identity of 27% with group A GraS but 58% with group A BraS. *S. pettenkoferi* BraR has a mean sequence identity of 47% with GraR and 73% with BraR from group A. These values support the *S. pettenkoferi* TCS is a BraSR orthologue and its singular association raises questions about the evolution of BraSR in group A staphylococci. Gene duplication of GraSR in a group A species, with subsequent sequence divergence over time to BraSR and spread throughout group A species by horizontal gene transfer, is tempting to suggest. *S. pettenkoferi* having BraSR but not GraSR presents a challenge to this paralogue hypothesis. We propose two possibilities; *S. pettenkoferi* may have suffered deletion of *graSR* following acquisition of *braSR*, or *S. pettenkoferi* never acquired *braSR*, but rather its TCS evolved from ancestral genes. Such a scenario would enable group A organisms to acquire *braSR* from *S. pettenkoferi* as an additional and sufficiently divergent TCS locus.

Regardless of the origins of both TCSs, the divergence between and within GraSR and BraSR likely reflect specificities for their ligands and selection driven by the niches to which the staphylococci are specialized.

### GraSR and BraSR-associated ABC transporters

Both GraSR and BraSR, as members of the BceS-like IM-HK family of TCS, are activated by AMP ligand bound to an associated ABC transporter (33). Given the important function of these TCS, the conservation of their associated transporter protein sequences was compared across the staphylococci.

VraFG is the GraSR-associated ABC transporter (34) and in the genomes encoding VraFG (absent from group B species and *S. pettenkoferi*) there is a high degree of shared protein sequence conservation. VraF has a mean sequence identity of 68% across the staphylococci examined (Table 1), with greatest conservation within species groups (group A, 79% identity; group B, 85.3% identity; group C, 96.8% identity). Shared sequence identity among the VraG proteins was 47.5%, with 88%, 65.2% and 61.9% identity within groups A, B and C, respectively. The BraDE ABC transporter associated with BraSR was identified in group A species and, similar to VraFG, revealed greater identity (68.4%) across BraD sequences compared with BraE (38.9%) protein sequences. Divergence within BraSR and GraSR-associated transporters has likely arisen from concurrent evolution of the ABC transporter specificities for AMPs.

### Experimental evolution of nisin resistance in *S. aureus*, *S. hominis* and *S. saprophyticus*

Previous studies demonstrated that selection by experimental evolution identified mutations conferring antimicrobial resistance in overarching regulators, notably SNPs in *braS* revealed roles for BraSR in nisin sensing and survival (7). Following our identified species restriction of BraSR to group A staphylococci, we adopted an experimental evolution strategy to interrogate the contributions of GraSR and BraSR TCS under selection for nisin resistance.

Strains of group A species, *S. aureus* and *S. hominis* plus group B *S. saprophyticus* were each serially passaged in triplicate cultures with increasing concentrations of nisin using a microtiter plate method, with an equivalent sodium citrate buffer control passaged in parallel. Stepwise increases in nisin MIC were observed for all strains tested with no obvious pattern in the rate of resistance acquisition between the species. After selection, both *S. aureus* 171 and *S. aureus* SH1000 strains exhibited ∼100-fold increases in nisin MIC, a greater fold increase in resistance than that observed by Blake *et al* (7), which may be due to experimental design differences. Selection of both *S. hominis* strains increased nisin MIC ∼25-fold, and *S. saprophyticus* strains CCM_883 and CCM_349 showed 80-fold and 5-fold increases, respectively. Multiple clones of *S. aureus* 171, *S. hominis* J31 and *S. saprophyticus* CCM883 were genome sequenced to identify sequence variants that potentially contributed to increased nisin MIC. T0 genomes were assembled and annotated, then reads from three pools (each comprising 5 independent clones) and one individual clone of each experimentally evolved species were aligned to their respective assembled genomes to identify sequence variants (SNPs, insertions/deletions) specific to nisin selection (Tables 4-6).

**Table 4:** Non-synonymous, homozygous SNPs from independent clone pools of staphylococci after nisin selection. Pools comprise 5 clones from each of three independent experiments. Nisin MICs of clones in each pool were confirmed to ensure they were similar.

| Gene ID (Prokka) | Protein ID | Pool | Position | Base change | Amino acid change | Allele frequency |
| --- | --- | --- | --- | --- | --- | --- |
| ***S. aureus* 171** |  |  |  |  |  |  |
| *walK* | WalK Sensor kinase | 1 | 17119 | A -> G | H<sub>364</sub>R | 1 |
| *gltB_1* | Glutamate synthase | 1 | 437361 | A -> T | Q<sub>797</sub>L | 1 |
| *rpoB* | DNA-directed RNA polymerase subunit beta | 1 | 520176 | C -> T | H<sub>506</sub>Y | 1 |
| *mraY* | Phospho-N-acetylmuramoyl-pentapeptide transferase | 1 | 1103071 | G -> A | V<sub>266</sub>I | 1 |
| *yhcF* | Transcriptional regulator | 1 | 1998279 | G -> A | G<sub>73</sub>R | 1 |
| hypothetical | Membrane protein | 2 | 615003 | C -> T | Q<sub>57</sub>* | 0.35 |
| *rpoC* | DNA-directed RNA polymerase subunit beta | 3 | 523690 | G -> T | A<sub>448</sub>S | 0.99 |
| *femB* | FemB | 3 | 1323450 | G -> A | R<sub>215</sub>H | 0.63 |
| phage terminase | Terminase | 3 | 1491745 | G -> C | G<sub>240</sub>A | 1 |
| *greA* | GreA | 3 | 1625376 | A -> T | L<sub>76</sub>* | 0.37 |
| *braS* | BraS sensor histidine kinase | 3 | 2627088 | C -> T | T<sub>175</sub>I | 1 |
| ***S. hominis* J31** |  |  |  |  |  |  |
| *rpoB* | DNA-directed RNA polymerase subunit beta | 1 | 41317 | G -> T | D<sub>1046</sub>Y | 0.79 |
| *graS* | GraS sensor histidine kinase | 2 | 147503 | C -> T | S<sub>120</sub>L | 0.42 |
| *ftsH* | FtsH Zinc metalloprotease | 2 | 2172470 | C -> A | D<sub>171</sub>E | 1 |
| *gmk* | Guanylate Kinase | 3 | 552630 | G -> A | R<sub>135</sub>H | 1 |
| *yhcF* | Transcriptional regulator | 3 | 1212777 | C -> T | Q<sub>47</sub>* | 1 |
| ***S. saprophyticus* 883** |  |  |  |  |  |  |
| *graS* | GraS sensor histidine kinase | 1 | 2068987 | G -> T | G<sub>209</sub>C | 1 |
| *codY* | CodY regulator | 2,3 | 1537681 | C -> A | L<sub>79</sub>F | 1,1 |
| *pitA* | Phosphate transporter | 2,3 | 2066724 | C -> T | A<sub>195</sub>V | 0.99,1 |
| *graS* | GraS sensor histidine kinase | 2,3 | 2069134 | C -> A | R<sub>160</sub>S | 1,1 |
| marR family | MarR regulator | 2,3 | 2136907 | C -> T | T<sub>62</sub>I | 1,1 |

**Table 5:** Non-synonymous, homozygous SNPs from single clones of *S. aureus, S. hominis* and *S. saprophyticus* after nisin selection. Pools comprise 5 clones from each of three independent experiments. Nisin MICs of clones in each pool were confirmed to ensure they were similar.

| ***S. aureus* strain 171 (single clone from pool 1)** |  |  |  |  |
| --- | --- | --- | --- | --- |
| **Gene ID (Prokka)** | **Protein ID** | **Position** | **Base change** | **Amino acid change** |
| *walK* | WalK Sensor histidine kinase | 17119 | A -> G | H<sub>364</sub>R |
| *gltB_1* | Glutamate synthase | 437361 | A -> T | Q<sub>797</sub>L |
| *rpoB* | DNA-directed RNA polymerase subunit beta | 520176 | C -> T | H<sub>506</sub>Y |
| *mraY* | Phospho-N-acetylmuramoyl-pentapeptide transferase | 1103071 | G -> A | V<sub>266</sub>I |
| *msrR* | Regulatory protein MsrR | 1309845 | G -> T | E<sub>181</sub>* |
| *yurK* | Transcriptional regulator | 1998279 | G -> A | G<sub>73</sub>R |
| ***S. hominis* strain J31 (single clone from pool 2)** |  |  |  |  |
| *ftsH* | Zinc metalloprotease FtsH | 2172470 | C -> A | D<sub>171</sub>E |
| ***S. saprophyticus* strain 883 (single clone from pool 1)** |  |  |  |  |
| *graS* | GraS sensor histidine kinase | 2068987 | G -> T | G<sub>209</sub>C |

**Table 6:** INDELs from nisin selection pools and single clones of *S. aureus, S. hominis* and *S. saprophyticus*. Insertion/deletion sequence change effects: frameshift (fs); upstream variant (uv); downstream variant (dv); deletion (del); insertion (ins). INDELs marked § are predicted to have a major consequence by SnpEFF.

| Source | Gene ID (Prokka) | Protein ID | Location | Base change | Effect |
| --- | --- | --- | --- | --- | --- |
| ***S. aureus* 171** |  |  |  |  |  |
| Single, Pool 3 | *lpl2_2* | Lipoprotein | 403168 | 195_196 insGG | I66fs § |
| Single, Pool 1,3 | *sdrE_1* | MSCRAMM family adhesin | 554844 | 2672_2673 insC | K891fs § |
| Single |  |  | 1990375 | -1_-1 insCC | uv |
| Single, Pool 2,3 |  |  | 2052262 | *1530_*1530 delTG | fs |
| Single, Pool 3 | *deoC2* | Deoxyribose-phosphate aldolase 2 | 2126552 | 450_451 insAG | K151fs § |
| Single, Pool 1 | hypothetical | Hypothetical protein | 2471437 | 116_117 insT | E40fs |
| Single, Pool 3 | *fnbA_2* | Fibronectin binding protein | 2490720 | 1763_1764 delCG | S588fs § |
| Pool 1 | hypothetical | Hypothetical protein | 1556011 | 201 delT | S67fs § |
| Pool 2 |  |  | 1337112 | -1_-1 insATG |  |
| Pool 2,3 | hypothetical | transposase | 1337623 | 209_211 delAAG | E70del |
| Pool 2 | hypothetical | transposase | 1819467 | 1047_1048 insC | *350fs § |
| Pool 2 | *leuA_2* |  | 2052046 | *1530 delC | dv |
| Pool 2 | hypothetical | hypothetical | 2471438 | 115_116 insATA | P39del, insHT |
| Pool 3 | *lpl2_1* | hypothetical | 402338 | 212_213 insCT | Q71fs § |
| Pool 3 | *sdrD_1* | MSCRAMM family adhesin | 550856 | 3237_3238 insC | M1080fs § |
| Pool 3 | hypothetical |  | 555082 | -1_-1 insG |  |
| Pool 3 | hypothetical | LPXTG surface protein | 2480756 | 216 delA | T72fs § |
| ***S. hominis* J31** |  |  |  |  |  |
| Single, Pool 2 | *ftsH* | Zinc metalloprotease | 2172279 | 325 delA | S109fs § |
| Pool 1 | *relA* | GTP Pyrophosphokinase | 951655 | 764 delA | Q255fs § |
| Pool 2 | *ssaA2_2* | CHAP domain containing protein | 1437614 | 575_578 delGTTA | G192fs § |
| ***S. saprophyticus* 883** |  |  |  |  |  |
| Single, Pool 1,2,3 | *ptsG* | PTS alpha-glucoside transporter subunit IIBC | 604054 | -1_-1 insAA | uv |

### Nisin-selected SNPs in staphylococci

Experimental evolution of *S. saprophyticus* identified a SNP in *graS* (GraS: A_160_S; table 4) that was present in two clone pools, and SNP *graS* G_209_C in a third pool. A single clone sequenced from the latter pool identified only one SNP in *graS* (GraS: G_209_C) and an upstream variant associated with *ptsG* (table 4). These data provide support for GraSR contributing to nisin resistance in *S. saprophyticus* given the absence of the BraSR TCS in this group B *Staphylococcus* species. Aside from TCS, other regulators may contribute to the nisin response in *S. saprophyticus* as evidenced by an identical SNP identified in two separate nisin resistance selections (pools 2 and 3) corresponding to a T_62_I change in an uncharacterized MarR transcriptional repressor.

In both *S. aureus* and *S. hominis* there are multiple pathways to high-level nisin resistance. Each species revealed SNPs in TCS systems, but these differed across the parallel selection experiments (Table 4-6). In *S. aureus*, a non-synonymous SNP in *braS* (BraS: T_175_I) was present in 100% of reads from one sequenced pool, differing from previous work that identified a discrete *braS* SNP (BraS: A_208_E) (7). Evidence for a second TCS contributing to nisin resistance arose from a *walK* non-synonymous SNP (WalK: H_364_R) within the diverse and flexible signal sensing PAS domain of WalK in *S. aureus* (35). WalKR is essential and functions to maintain cell wall metabolism (36) and SNPs in this TCS contribute to vancomycin and daptomycin resistance due to cell wall-thickening (37). Should this cell wall phenotype be associated with the H_364_R WalK variant it could similarly limit nisin interaction with its lipid II target to abrogate pore formation. A large overlap was reported between the WalKR and GraSR regulatory networks in *S. aureus* (6).

In *S. hominis*, a *graS* SNP (GraS: S_120_L) was present in 2 clones of sequence pool 2 and no SNPs or other sequence variants were identified in *braSR* (Table 4-6). *S. hominis* has both *braSR* and *graSR* loci and therefore it is intriguing nisin resistance selection resulted in SNPs in a different TCS to *S. aureus* despite encoding both, potentially reflecting differences in their contribution across group A staphylococci. A further transcriptional regulator might contribute to nisin resistance in both *S. aureus* and *S. hominis*, where the uncharacterized *yhcF* revealed SNPs producing G_73_R and N_47_*, respectively; the presence of SNPs in *yhcF* of both species supports a role for this regulator. The YhcF transcriptional regulator proteins of *S. aureus* and *S. hominis* have 75% similarity and their cognate genes are adjacent to an ABC transporter locus with potential specificity for GlcNAc, which might catalyze recycling of cell wall substrates from nisin damage. The role of this operon is currently being investigated.

In summary, we have identified differential encoding and diversity of antimicrobial resistance regulators and their associated transporters across the staphylococci. Our previous studies of the nasal microbiome correlated cumulative antimicrobial production with community structure, limitation of invasion and *S. aureus* exclusion (38, 39, 40). Further dissection of antimicrobial sensing and discrimination via the TCS systems BraSR and GraSR combined with analysis of their associated transport specificities will provide information that can be layered with niche-relevant antimicrobial activities from competing species. Such analyses are now emerging and will provide a more holistic determination of *Staphylococcus* ecology.

## Methods

### *Staphylococcus* orthologous gene content

Representative genomes of 21 different *Staphylococcus* species available at the time of analysis (Table 1) were either sequenced (see later section) or retrieved from the NCBI FTP repository (ftp://ftp.ncbi.nlm.nih.gov/). Complete genomes were used where possible. Draft genomes available as NCBI scaffolds were reordered against an appropriate reference using a bespoke perl script. Genomes were annotated using PROKKA (version 1.5.2) (41) to ensure consistent gene calling and annotation. OrthoMCL (version 1.4) was used to cluster orthologous proteins (42), with input parameters, e-value cut-off: 1e-5, percentage identity cut-off: 30, percentage match cut off: 20. A bespoke python script was used to create a table describing the presence or absence of each OrthoMCL cluster within every genome. These data were converted to a matrix for analysis in the statistical package R and a heatmap was generated from the matrix. To control for gross strain-specific effects on the heat map (and thus OrthoMCL clusters), this step was repeated by substituting with alternative strains (Table S1) and all permutations were analyzed in subsequent steps of the analysis.

### Drivers of OrthoMCL group formation

The R library, Random Forest (version 4.6-7) (43) was used to investigate the genetic inputs directing classification of the species into their OrthoMCL groups. A presence/absence table of each of the orthologous groups obtained from the USA300 permutation of the OrthoMCL analysis was generated using a bespoke python script and used as the input data for the Random Forest algorithm.

The data was split into a test and training data set with both sets including equal proportions of group A species. The optimum value for mtry was found to be 66 using the tuneRF function (ntree=1001, stepFactor=1.5, improve=0.001). These mtry and ntree parameters resulted in a model with an out of bag error rate of 9.09%.

Data output was summarized using the variable importance plot function and the numeric mean decrease in accuracy (MDA) resulting from the permutation of each variable was obtained through the importance function; these data were used as the measure of the importance of each variable. The maximum MDA in this analysis was 3.3. Clusters were mapped back to the genome and the annotation of protein sequence for a species representative of each cluster was retrieved. Protein sequences of clusters identified as important were retrieved and their annotations curated and verified against published annotations. In addition, outputs were generated by substituting strains of species in the analysis to compare conservation of identified clusters between the variable importance plots. Sequences of protein clusters from the single species representative in Table 2 and identified by Random Forest output are listed in Supplementary Files S2-3. Protein sequences were retrieved from their respective genomes and alignments were performed using ClustalW2 (version 2.1).

### Minimum inhibitory concentration assay

Nisin (Sigma-Aldrich Company Ltd, UK) was prepared as a 20 mg mL^−1^ solution in 10 mM sodium citrate (Sigma-Aldrich Company Ltd, UK) at pH 3 and stored at 4 °C. MIC assay used microtiter plates with doubling dilutions of nisin in BHI (Thermo Scientific) inoculated 1 in 2 with 100 μL bacterial suspension adjusted to OD_600_ 0.2 ± 0.005. The lowest concentration with an optical density ≤ to that of the initial optical density was taken as the minimum inhibitory concentration (MIC).

### Selection for nisin resistance

Experimental evolution was performed by serial passage in broth containing doubling dilutions of nisin in triplicate wells of a microtiter plate. For selection of *S. aureus* and *S. saprophyticus*, the maximal assay concentration of nisin was 5 mg mL^−1^ and for *S. hominis* 50 μg mL^−1^. Control selection experiments with equivalent sodium citrate concentrations were performed in parallel. Experiments were initiated with inoculation of bacteria to OD_600_ =0.2 for the first passage and plates were incubated static at 37 °C. Bacteria growing at the highest concentration of nisin after 24-48 h were passaged forward to the next plate; subsequent passages were inoculated with a 1:1000 dilution of culture. Serial passage was continued until growth occurred at the maximal nisin concentration (for strains *S. saprophyticus* =10 mg ml^−1^, *S. aureus* =10 mg ml^−1^ and *S. hominis* =250 μg ml^−1^) or for a period of 12 days. All passaged cultures were collected and stored at −80°C in 20% (v/v) glycerol (Fisher Scientific) after each passage and the T_0_ time point served as comparator strain.

Colonies were randomly selected for sequencing after plating from independent biological replicate cultures that had reached an equivalent maximum level of nisin resistance. Clones from each repeat were selected and cultured in 10 mL of BHI at 37 °C with shaking at 200 rpm overnight. Increased MICs were confirmed by using the MIC assay described above at the highest nisin concentrations. Selection was performed for a corresponding citrate control time point for each of the three species.

### DNA extraction, library preparation and sequencing

Cells were harvested from overnight culture and lysed in buffer containing 12.5 μg ml^−1^ lysostaphin (Sigma-Aldrich) and 10 U mutanolysin (Sigma-Aldrich). DNA was purified using a DNeasy Blood and Tissue Kit (Qiagen). DNA (30 ng) from each of five selected clones was pooled to make Illumina Truseq DNA libraries with an insert size of 350 bp. In addition to three separate clone pools, a single clone was selected for sequencing from the clones used to constitute the pools. Single clones were selected on the basis of the highest DNA quality. The single clones and the T_0_ isolates were also sequenced using Illumina Truseq nano DNA libraries with 350 bp inserts.

### Identification of SNPs and INDELS

T_0_ comparator strains were assembled using VelvetOptimiser (version 2.2.5; Victoria Bioinformatics Consortium) with Kmer sizes from 19 to 99 and Velvet version 1.2.06 (45). Annotation was carried out using PROKKA version 1.5.2 (Seemann 2014). The PacBio assembly of *S. hominis* strain J31 (Accession FBVO01000000) (46) was used as the comparator assembly for this strain. Good quality filtered reads from experimentally evolved pools and single clones were aligned to respective comparator strains using the BWA (version 0.5.9-r16) (47) packages aln and sampe, and also using BWA (version 0.7.5a-r405) mem package. SAM files were converted to bcf (binary variant call) files with samtools for SNP calling using the mpileup package. The bcf output file from mpileup was then converted to vcf (variant call format) files and quality filtered. For SNPs, only this quality filtered vcf file from the pooled clones, along with mpileup output without base data, were used to further filter the SNPs to include only those present in 33.33% of reads, which equates to the SNP being present in more than one clone. To reduce falsely called SNPs, SNPs not called from both alignments (from either BWA aln and sampe or BWA mem) were removed from the data set, as recommended by (48). SNPs called in the control data and evolved isolates were filtered from the data.

## Data accession

Genomes resulting from this work can be retrieved from the ENA database at EMBL-EBI (http://www.ebi.ac.uk/ena/data/view) under the bioproject accession PRJEB22856, including data from experimental evolution of *S. aureus* 171; Parental *S. aureus* 171 data accession: LT963437. Individual genome assembly accessions used in Figure 1 are listed in Table 1 and Supplementary File S1.

## Acknowledgements

We are grateful to Dr Miriam Korte-Berwanger, University of Bochum and Prof Ross Fitzgerald, University of Edinburgh for kindly providing *Staphylococcus* strains used in this study.

---

**File S1. *Staphylococcus* species and strains used as substitutes in OrthoMCL analyses.**

**File S2. Species group A, present and absent cluster protein sequences.** Sequences represent clusters listed in Table 2.

**File S3. Species group B orthologue sequence.** Sequence of putative squalene synthase cluster of *S. xylosus*.

